# Synchronization of the prefrontal cortex with the hippocampus and posterior parietal cortex is navigation strategy-dependent during spatial learning

**DOI:** 10.1101/2024.05.15.594336

**Authors:** F. García, M-J. Torres, L. Chacana-Véliz, N. Espinosa, W. El-Deredy, P. Fuentealba, I. Negrón-Oyarzo

## Abstract

During goal-directed spatial learning, subjects progressively change their navigation strategies to increase their navigation efficiency, an operation supported by the medial prefrontal cortex (mPFC). However, how the mPFC may integrate relevant information in a wider memory networks involving the hippocampus (HPC) and the posterior parietal cortex (PPC) is poorly understood. We recorded local-field potential and neuronal firing simultaneously from the mPFC, HPC and PPC in mice subjected to spatial memory acquisition in the Barnes maze. During navigation trials, animals demonstrated two consecutive behavioral stages: searching and exploration. Throughout training, mice gradually switched from less efficient (non-spatial) to more efficient (spatial) goal-oriented strategies exclusively during the searching stage. 4-Hz and theta (6-12 Hz) oscillations were detected during spatial navigation in the three recorded areas associated with episodes of immobility and locomotion, respectively. The entrainment of prefrontal gamma oscillations (60-100 Hz) by hippocampal and parietal 4-Hz and theta oscillations, as well as the incidence of prefrontal gamma, was higher when mice implemented spatial strategies during the searching stage. Interestingly, 4-Hz and theta from HPC and PPC also synchronized the spike-timing of prefrontal neurons, which was maximum during spatial strategies in the searching stage. Finally, neurons recorded in the mPFC increased their task stage firing selectivity when they used spatial strategy. Altogether, these results provide evidence for the neural mechanisms underlying the prefrontal large-scale coordination with distributed neural networks during spatial learning.

## 2. INTRODUCTION

During goal-directed spatial memory acquisition, subjects progressively implement navigation strategies of increasing efficiency to reach the goal (DiMattia & Kesner, 1988; Harrison et al., 2006; Negrón-Oyarzo et al., 2018; Ruediger et al., 2012). Considering current learning theories, this phenomenon may reflect the execution of the active guidance of behavior product of the cumulated experience, for which the prefrontal cortex (PFC) plays a critical role (Eichenbaum, 2017; Preston & Eichenbaum, 2013; Schlichting & Preston, 2015). Interestingly, strategy progression during spatial learning is mediated by the medial PFC (mPFC) (de Bruin et al., 1994; Kesner et al., 1989; Kolb et al., 1994; Patai & Spiers, 2021). Indeed, during spatial tasks, firing patterns in the mPFC represent several cognitive features relevant for strategy progression, such as strategy switching and implementation of efficient navigation strategies (Malagon-Vina et al., 2018; Negrón-Oyarzo et al., 2018; Powell & Redish, 2016; Rich & Shapiro, 2009). This suggests a critical role for the mPFC in executive aspects of spatial learning. For this, the mPFC may integrate information represented across distributed brain areas (Euston et al., 2012). Information concerning spatial-temporal sequences and active guidance of the body through the visual space is crucial for spatial learning (Save & Poucet, 2000; Whitlock et al., 2008), which are represented in the hippocampus (HPC) and posterior parietal cortex (PPC), respectively (Alexander et al., 2022; Buzsáki & Tingley, 2018; Minderer et al., 2019; Nitz, 2006). These structures are required for spatial learning (DiMattia & Kesner, 1988; Kesner et al., 1989; Kolb et al., 1994) and are anatomically connected with the mPFC (Jay & Witter, 1991; Kolb & Walkey, 1987; Swanson, 1981; Zingg et al., 2014). Therefore, the mPFC may integrate relevant information from the HPC and PPC for strategy progression during learning. However, the mechanism underlying the prefrontal coordination with HPC and PPC during spatial learning is poorly understood.

It has been proposed that transient synchronization of local and distributed neural activity patterns may support integration of information required for cognitive operations (Engel et al., 2001; Fell & Axmacher, 2011; Fries, 2005; Varela et al., 2001). Low-frequency oscillations, such as theta (6-12 Hz) and 4-Hz oscillations, synchronize gamma activity (30-100 Hz) and firing patterns in the mPFC (Fujisawa & Buzsáki, 2011; Sirota et al., 2008; Tamura et al., 2017). Simultaneously, gamma oscillations, which represent local neural operations underlying information processing (Fernandez-Ruiz et al., 2023), coordinate firing of neural populations into particular computations, supporting the formation of task-relevant firing patterns in the mPFC (Fujisawa & Buzsáki, 2011). Thus, low-frequency oscillations, though long-range entrainment of gamma activity and firing patterns, may provide windows of efficient communication between distant neural networks, promoting large-scale information transfer for the implementation of task-related neural operations in the mPFC (Buzsáki & Wang, 2012; Canolty & Knight, 2010; Griffiths & Jensen, 2023; Hyafil et al., 2015). We propose that this mechanism may support strategy progression during spatial learning.

In this study, we performed simultaneous large-scale recordings of neuronal activity and local field potential (LFP) in the mPFC, the HPC, and the PPC of mice during a spatial memory acquisition task. Along navigation trials, we identified searching and exploratory task stages, in which strategy progression was observed during searching but not during exploration. Theta and 4-Hz oscillations were evident in the three recorded areas in relationship with locomotor activity. Importantly, gamma oscillations and neuronal firing in the mPFC were coordinated by hippocampal and parietal theta and 4-Hz oscillations according to task stages in a strategy progression-dependent manner. Lastly, firing patterns in the mPFC were dynamically modulated by strategy progression. Altogether, these results provide evidence for the neural mechanisms underlying the large-scale coordination of distributed neural networks during spatial learning.

## 3. RESULTS

### 3.1. Identification of task stages and strategy progression during spatial learning

Adult mice (n = 12) were chronically implanted with microelectrodes simultaneously in the mPFC, HPC and PPC and were subjected to training in the Barnes maze (**Fig. 1a**). In this behavioral paradigm, animals must escape from an aversive (elevated and illuminated) arena in which an escape hole is located in a fixed spatial location across all training trials (**Fig. 1a**); thus, a goal-directed spatial memory is formed across navigation trials. Representative occupancy colorplots depicted in **supplementary Fig. 1a** shows a progressive decrease in the path length to the goal and an increased occupancy near the goal across training days. Escape latency and the number of errors were significantly reduced across training days (latency: P = 0.026; **Fig. 1b**; errors: P = 0.025; one-way ANOVA; **supplementary Fig. 1b**). These results were not related to modification of locomotor activity, as no significant changes in mean and maximum running speed were observed across training days (mean speed: P = 0.719; max. speed: P = 0.951; one-way ANOVA; **supplementary Fig. 1c, d**).

**Figure 1.**
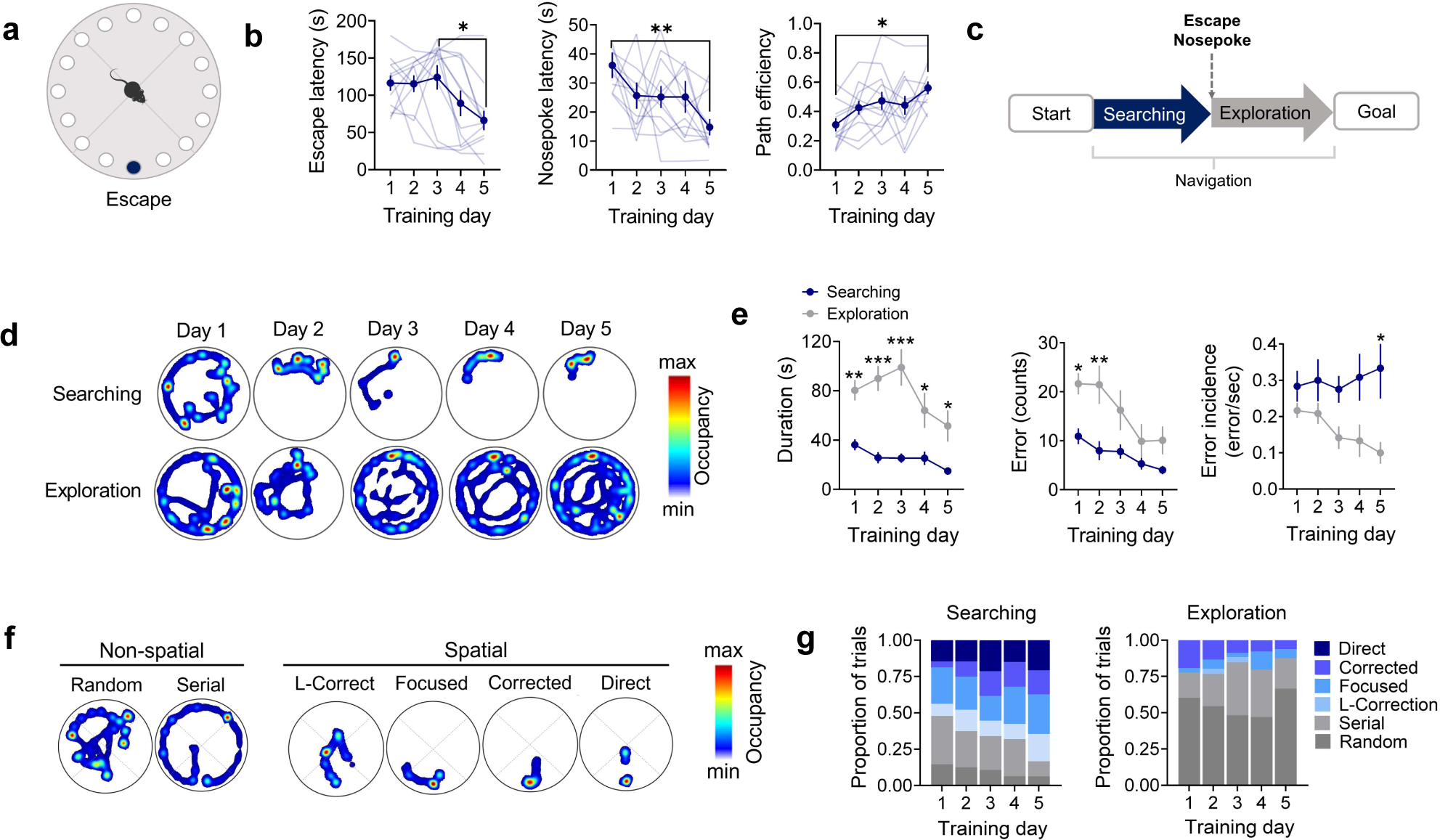
Behavioral performance during spatial learning in the Barnes maze. **a)** Schematic diagram of the Barnes maze. **b)** Escape latency (left), goal-nose poke latency (middle), and path efficiency to nose poke in the escape hole (right) across acquisition days in the Barnes maze. Clear blue is data of individual mice; solid blue indicates the mean ± SEM. *: P < 0.05; **: P < 0.01; Tukey’s multiple comparisons test after one-way ANOVA. **c)** Schematic diagram of the criterion for distinction between searching and exploration task stages. **d)** Example occupancy colorplots in the Barnes maze during the searching and exploration task stages of the same trial for each acquisition day. **e)** Comparison of average duration (left), error counts (middle), and error incidence across training days between searching and exploration task stages. Data are presented as mean ± SEM. *: P < 0.05; **: P < 0.01; Sidak’s multiple comparisons test after two-way ANOVA. **f)** Example occupancy colorplots of the navigation strategies used to find the goal in the Barnes maze. Strategies were classified as non-spatial or spatial if animals were directed toward the goal quadrant. **g)** Normalized distribution of navigation strategies implemented across training days during the searching and exploration task stages.

We observed that mice do not necessarily enter the escape hole once they find it. Indeed, it has been considered that the first nose poke into the escape hole is a more reliable indicator of spatial learning (Harrison et al., 2006). We found that escape-nose poke latency and error counts before escape-nose poke significantly decreased across training days (latency: P = 0.023; errors: P = 0.020; one-way ANOVA; **Fig. 1b and supplementary Fig. 1e**, respectively), without changes on mean and maximum speed (mean speed: P = 0.635; maximum speed: P = 0.221; one-way ANOVA; **supplementary Fig. 1f, g**). Importantly, path efficiency to escape-nose poke increased significantly along training days (P = 0.033; one-way ANOVA; **Fig. 1b**), suggesting a progressive increase in the proficiency to find the escape. A detailed inspection of the trajectories suggests that after finding the escape, mice implemented a different behavior that was not necessarily related to finding the escape but related to the exploration of the unfamiliar environment. To evaluate this possibility, we divided each navigation trial into two task stages with respect to the first encounter with the escape hole: before the first nose poke in the escape hole (which we called the searching stage) and after the first nose poke in the escape hole (exploration stage; **Fig. 1c**). Representative occupancy plots for both task stages of the same navigation trial across training days are shown in **Fig. 1d**. We found that duration, error counts, and distance traveled across training days were lower in the searching stage compared to exploration (P < 0.0001; two-way ANOVA; **Fig. 1e** and **supplementary Fig. 1h**). We found no significant differences in mean or maximum speed between task stages (mean speed: P = 0.3860; max speed: P = 0.617; two-way ANOVA; **supplementary Fig. 1i, j**). Thus, mice spent most of the navigation time exploring the arena after they found the escape hole, even when they already knew its location. Occupancy plots in **Fig. 1d** also show that navigation was progressively concentrated near the goal in the searching stage, whereas mice navigated in a larger area of the arena during exploration. Indeed, path sparsity, which estimates the spatial distribution of the trajectory in the maze, was higher in the exploration stage compared to searching (P = 0.018; t-test; **supplementary Fig. 1k**). Finally, we reasoned that the motivation to find the escape could be reflected as continuous attempts to find escape across training days, which could be highest during searching compared to exploration. To test this, we calculated the error incidence, i.e., the number of error counts per second for each stage across training days. Error incidence was highest in searching (P < 0.001; two-way ANOVA; **Fig. 1e**). Indeed, error incidence remained almost persistent across training days during the searching stage, consistent with the motivation to find the escape. Contrarily, error incidence in the exploration stage decreased across training days, suggesting a decreased motivation to find the escape (**Fig. 1e**).

We then evaluated the progression of the navigation strategies across training. Given that escape-directed navigation was implemented during the searching stage, we first evaluated strategy progression in this stage. To this aim, navigation paths were qualitatively classified according to the Barnes maze Unbiased Strategy classification algorithm (BUNS; (Illouz et al., 2016)) which identifies six different navigation strategies: random, serial, focused search, long-correction, corrected and direct. Representative occupancy plots for each navigation strategy are shown in **Fig. 1f**. We found significant differences across navigation strategies in path efficiency to goal-nose poke, path sparsity, latency to goal-nose poke, and number of errors (P < 0.0001; one-way ANOVA; **supplementary Fig. 1l-o**), with no significant differences in mean speed (P = 0.454; one-way ANOVA; **supplementary Fig. 1p**). This suggests that navigation strategies were related to changes in the trajectory but not in the velocity, which strongly impacts path proficiency and behavioral performance during the searching stage. Importantly, less efficient strategies were mostly implemented early during training, whereas those more efficient were mainly evident at the end of training, with an apparent continual progression from less-to-more efficient strategies across training days (**Fig. 1g**); the significant increase of the BUNS-cognitive score across training days agrees with this observation (P = 0.045; one-way ANOVA; **supplementary Fig. 1q**). Finally, if the exploration stage is not directed toward finding the goal, then strategy progression would be absent during the exploration stage. Occupancy plots in **Fig. 1d** show that random navigation was principally observed during exploration (nearly 50% of trials), with no apparent strategy progression across training days. Indeed, the cognitive score did not change across training days in the exploration stage (P = 0.609; one-way ANOVA; **supplementary Fig. 1r**). Overall, these data show that two behavioral stages were clearly distinguishable during navigation: a first searching stage directed to finding the escape, in which goal-directed strategy progression was evidenced, and a consecutive exploratory but non-goal-directed stage with no evident strategy progression.

### 3.2. 4-Hz and theta oscillations emerged in the mPFC, HPC and PPC during spatial learning

We then asked if these behavioral findings were related to changes in neural activity patterns. LFP and neuronal firing were simultaneously recorded from the mPFC, HPC and PPC during all training sessions. Examples of Nissl-stained sections showing the position of recording electrodes are shown in **Fig. 2a**. Good-quality recordings were obtained from 10 of 12 recorded mice (20 trials per animal, a total of 200 recording sessions). As observed in the raw LFP recording and spectrograms in **Fig. 2b**, 4-Hz and theta oscillation (6–12 Hz) were evident in the three recorded areas. The comparison of the representative instant energy suggests that these oscillations emerge segregated (**Fig. 2c** and **supplementary Fig. 2a**), which seems to be related to locomotion (**Fig. 2c**). To evaluate this possibility, we identified moments of locomotion and immobility (running speed above and below 4 cm/s, respectively) during the searching and exploration stages and compared the z-scored spectral power for each oscillation. We found that the spectral power of 4-Hz oscillation was higher during immobility compared to mobility in the three recorded areas (mPFC: P = 0.044; HPC: 0.040; PPC: P = 0.048; paired t-student test; **Fig. 2d**). Contrarily, theta power was higher during mobility (mPFC: P = 0.041; HPC: 0.030; PPC: P = 0.005; paired t-student test; **Fig. 2d**). Task stages did not affect the power of both 4-Hz and theta oscillations in the three recorded areas (**supplementary Fig. 2b**).

**Figure 2.**
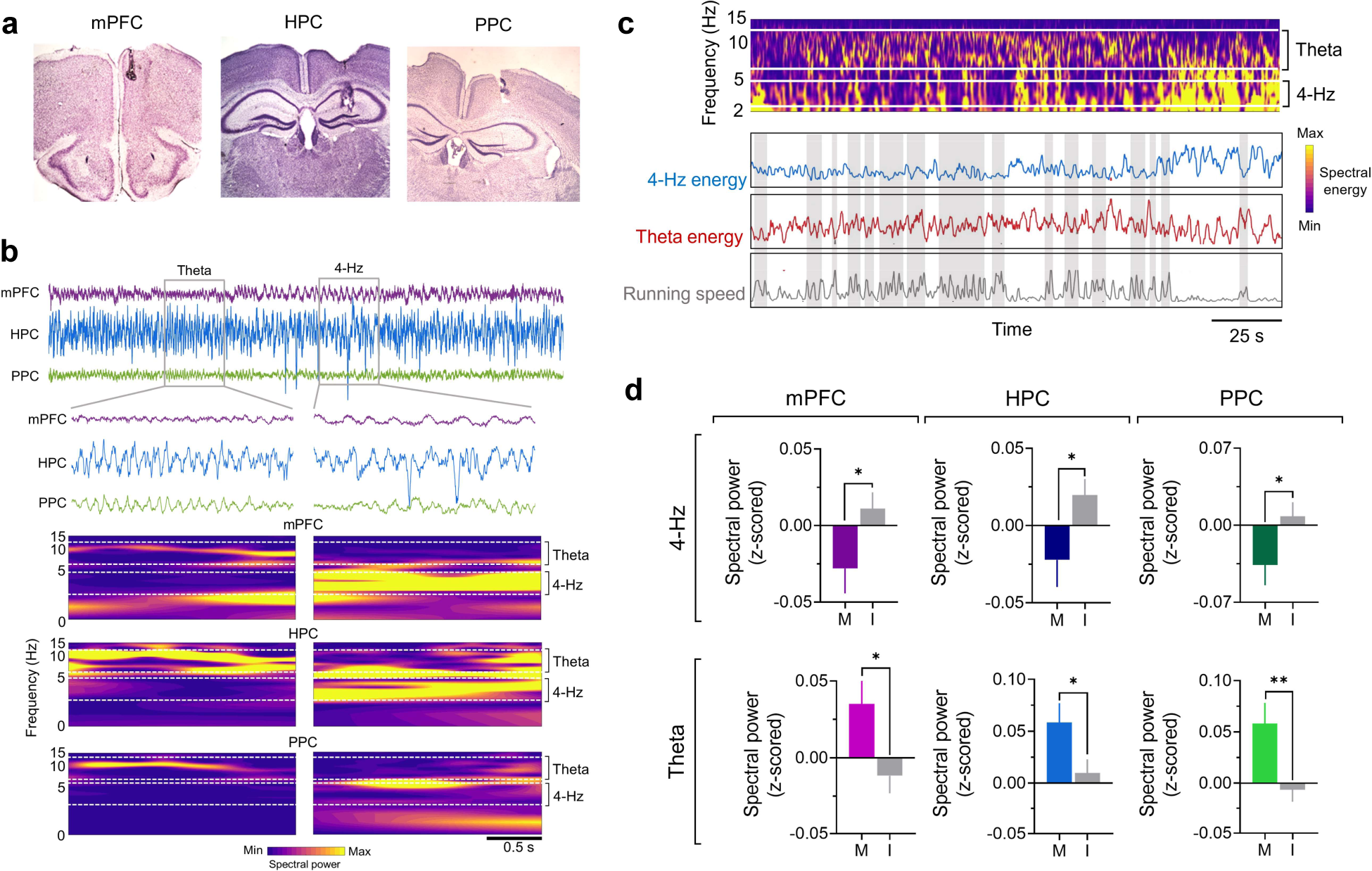
4-Hz and theta oscillations were detected in the mPFC, HPC, and PPC during spatial learning. **a)** Example of microphotography (magnification: 40x) showing the electrolytic mark in the mPFC (left), HPC (middle), and PPC (right). **b)** Upper panel: example of LFP recorded simultaneously from the mPFC, HPC, and PPC during training in the Barnes maze. Middle: magnification of segments of LFP recordings showing theta (left) and 4-Hz oscillations (right). Lower panel: spectrograms for LFP recordings shown in the middle panel. **c)** An example of the comparison of instantaneous energy at 4-Hz (upper panel) and theta frequency band (middle panel) of an LFP recorded from the mPFC aligned with the instantaneous speed. Gray areas are periods of locomotion (instantaneous speed over 4 cm/s). **d)** Bar chart of mean z-scored spectral power in the 4-Hz (upper) and theta-frequency band (lower) in the mPFC (left) HPC (middle) and PPC (right) in respect to mobility and immobility episodes. Data are presented as mean ± SEM. *: P < 0.05; Sidak’s multiple comparisons test after two-way ANOVA.

### 3.3. Hippocampal and parietal oscillations entrained prefrontal gamma oscillation according to task-stages and strategy progression during spatial learning

Given that both theta and 4-Hz oscillations are able to entrain gamma oscillations in the mPFC (Fujisawa & Buzsáki, 2011; Sirota et al., 2008; Tamura et al., 2017), we asked if hippocampal and parietal low-frequency oscillations entrained gamma oscillations in the mPFC during spatial learning. As shown in the raw and filtered recordings in **Fig. 3a**, the amplitude of prefrontal gamma activity seems to be coordinated by the phase of hippocampal and parietal 4-Hz and theta oscillations. Therefore, we computed the phase-amplitude coupling (PAC) of prefrontal gamma with respect to the phase of hippocampal and parietal oscillations (Tort et al., 2010). Representative comodulograms are shown in **supplementary Fig. 3a**. Hippocampal and parietal entrainment strength (measured as modulation index, MI) of prefrontal gamma was higher for 4-Hz compared to theta (P < 0.0001; Sidak’s multiple comparisons test after two-way ANOVA), with no significant differences between areas (4-Hz: P = 0.634; theta; P = 0.373; Sidak’s multiple comparisons test; **supplementary Fig. 3b**). The mean frequency for phases 4-Hz and theta was identical between HPC and PPC (4-Hz: P = 0.151; theta: P = 0.272, paired t-student test; **supplementary Fig. 3c**). Similarly, the mean frequency for prefrontal gamma amplitude modulated by 4-Hz and theta oscillations was identical between HPC and PPC (near 80 Hz; 4-Hz: P = 0.371; theta: P = 0.378; paired t-student test; **supplementary Fig. 3c**).

**Figure 3.**
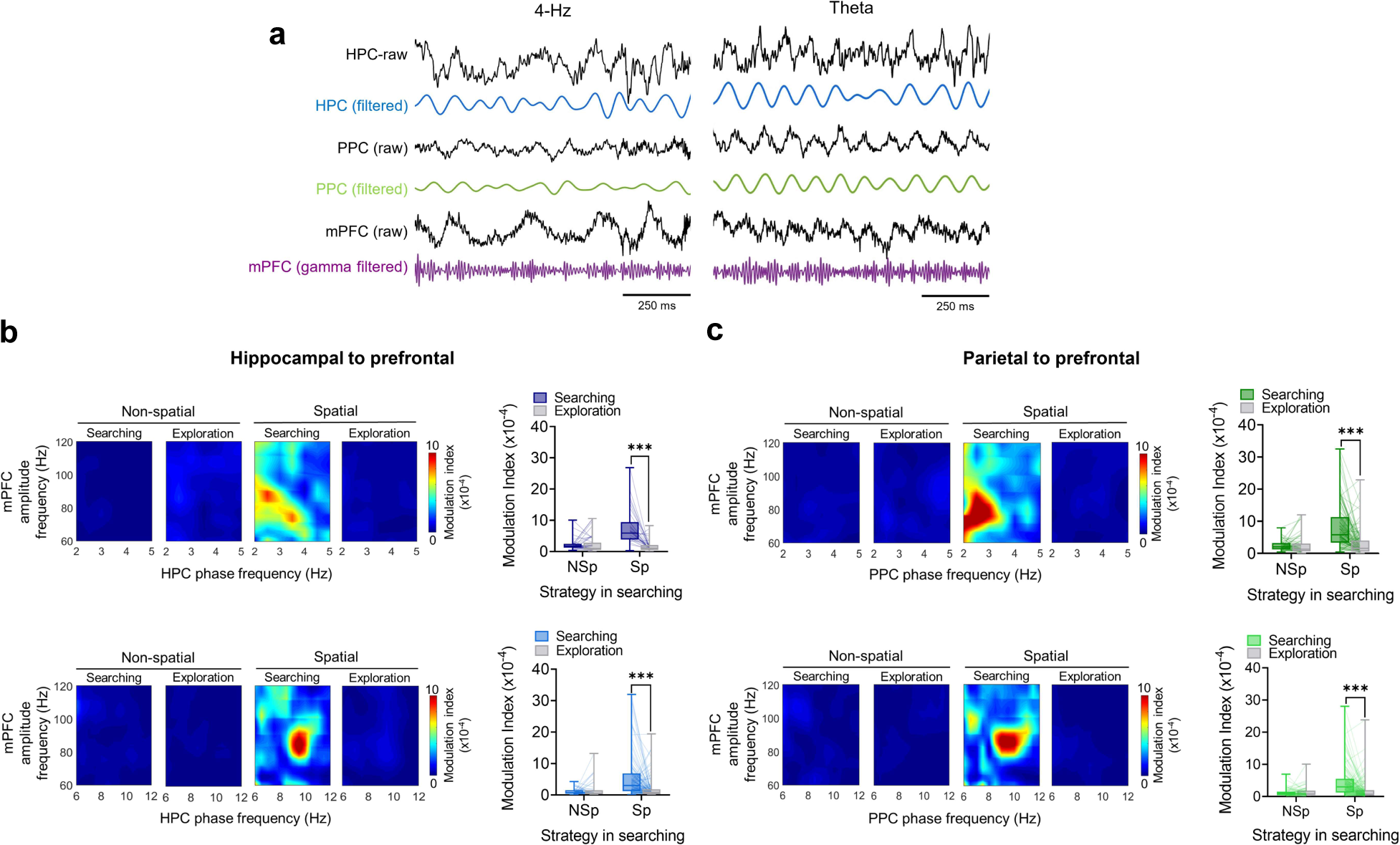
Prefrontal gamma oscillations are modulated by hippocampal and parietal 4-Hz and theta oscillations according to task requirements. **a)** Left: simultaneous LFP recordings from HPC (blue; raw and 4-Hz filtered), PPC (green; raw and 4-Hz filtered), and mPFC (purple; raw and gamma filtered). Right: simultaneous LFP recordings from HPC (blue; raw and theta filtered), PPC (green; raw and theta filtered), and mPFC (black; raw and gamma filtered). **b)** Upper: example of comodulograms (left) and box plot of modulation index (left) of prefrontal gamma oscillations with respect to the phase of hippocampal 4-Hz at different task stages and navigation strategies. Lower: same as upper panel, but for hippocampal theta.***: P < 0.001; Sidak’s multiple comparisons test after two-way ANOVA. **c)** Same as **b)**, but for prefrontal gamma modulation by parietal 4-Hz and theta oscillations. ***: P < 0.001; Sidak’s multiple comparisons test after two-way ANOVA. For all box plots, the middle, bottom, and top lines correspond to the median, lower, and upper quartiles, and the edges of the lower and upper whiskers correspond to the 5th and 95th percentiles.

We then asked if this long-range entrainment was modulated by navigation strategies at different task stages during spatial learning. To facilitate the analysis and data representation, we classified navigation strategies into two groups based on the criteria of movement toward the goal quadrant: random and serial strategies were considered “non-spatial,” whereas focused search, long-correction, corrected and direct were considered “spatial” (**Fig. 1f**) (Negrón-Oyarzo et al., 2018). Examples of comodulograms are shown in **Fig. 3b, c**. We found that 4-Hz and theta hippocampal entrainment of prefrontal gamma were highest in the searching stage respect to exploration during spatial strategies (4-Hz: P < 0.0001; theta: P < 0.0001; Sidak’s multiple comparisons test after two-way ANOVA; **Fig. 3b**) with no differences between stages in non-spatial strategies (4-Hz: P = 0.986; theta: P = 0.884; Sidak’s multiple comparisons test after two-way ANOVA; **Fig. 3b**). Also parietal entrainment of prefrontal gamma by 4-Hz and theta oscillations was highest in searching during spatial (4-Hz: P < 0.0001; theta: P < 0.0001; Sidak’s multiple comparisons test after two-way ANOVA; **Fig. 3c**), but not during non-spatial strategies (4-Hz: P = 0.986; theta: P = 0.666; Sidak’s multiple comparisons test; **Fig. 3c**). This highest entrainment observed in searching during spatial strategies was not attributed to the duration of the stages (**supplementary Fig. 3d**).

The increased long-range entrainment of prefrontal gamma may impact gamma activity in the mPFC. Therefore, we evaluated if gamma changed according to task stages and strategy progression during spatial training. Gamma oscillations appear as discrete oscillatory events (**Fig. 4a**); we identified these events and calculated the incidence of gamma events during searching and exploration stages for every navigation trial. We found an increase in the incidence of gamma events in the mPFC when animals used spatial strategies during searching compared to subsequent exploration (P = 0.0004; Sidak’s multiple comparisons test after two-way ANOVA; **Fig. 4b, c**), a difference not observed in non-spatial strategies (P = 0.080; Sidak’s multiple comparisons test after two-way ANOVA; **Fig. 4b, c**). No changes between stages and strategies were observed in the HPC and PPC (HPC; non-spatial: P = 0.609; spatial: P = 0.621; PPC; non-spatial: P = 0.993; spatial: P = 0.081; Sidak’s multiple comparisons test after two-way ANOVA; **Fig. 4b, c**).

**Figure 4.**
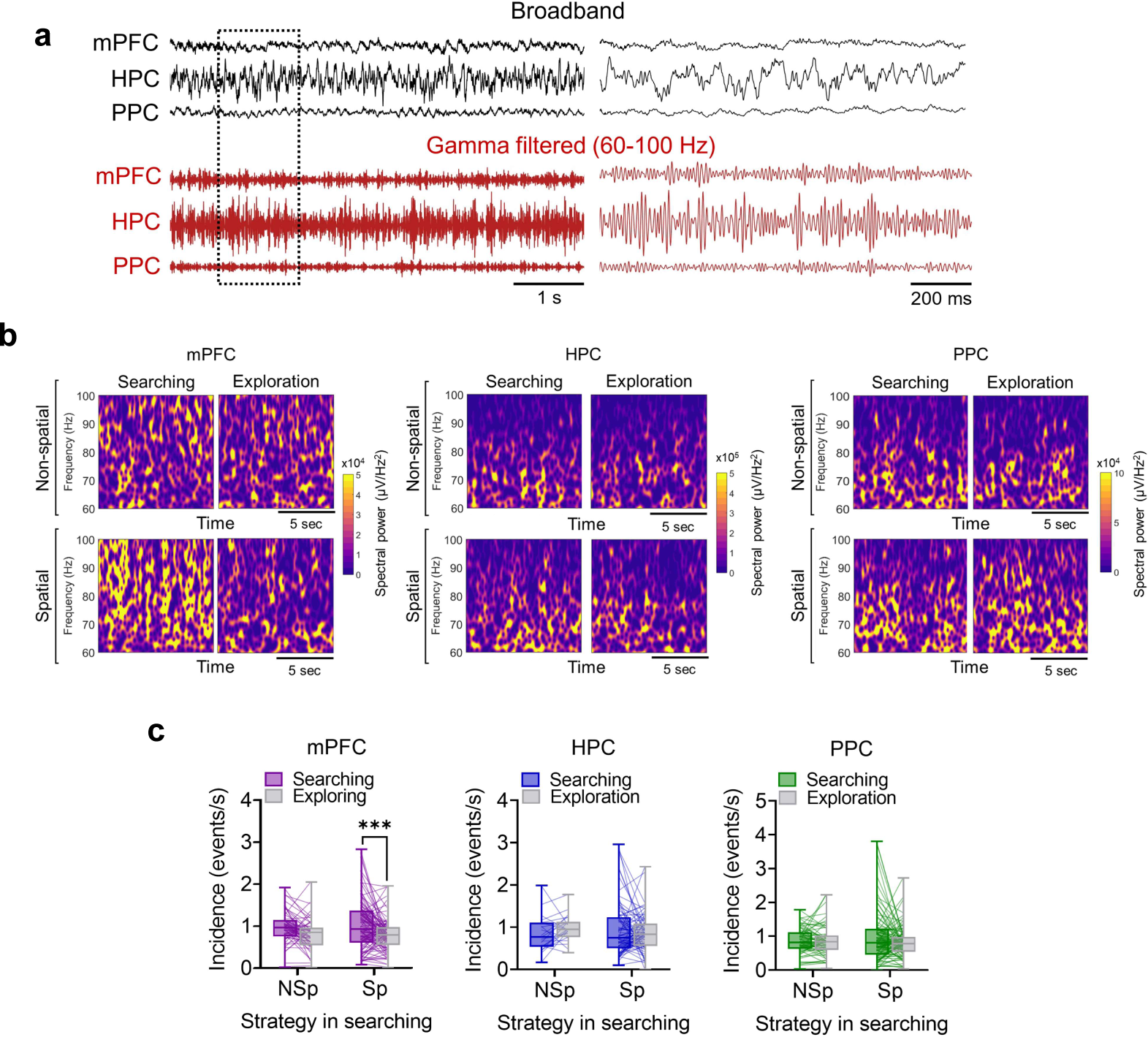
Gamma oscillations during spatial learning. **a)** Example of raw (upper, black traces) and gamma-filtered (lower, red traces) simultaneous LFP recordings from mPFC, HPC, and PPC. The panel on the right is a magnification of the dotted area from the left panel. **b)** Example of colorplot spectrograms of gamma oscillations of LFPs recorded from mPFC, HPC, and PPC during different task stages and navigation strategies. **c)** Box plots of the incidence of gamma events in the mPFC, HPC, and PPC with respect to task stages and to navigation strategies. ***: P < 0.001 Sidak’s multiple comparisons test after two-way ANOVA.

### 3.4. Local and large-scale oscillations synchronized the spiking of prefrontal neurons according to task stages and navigation strategies during spatial learning

Gamma oscillations are able to coordinate neural populations in cortical networks. Therefore, we asked if prefrontal gamma synchronized local neural firing and if this coordination was modulated by navigation strategies in different task stages. We obtained a total of 382 single unit recordings from the mPFC during training sessions. An example of simultaneous neurons modulated by local gamma oscillation is shown in the recording of **Fig. 5a**. To compare the strength of entrainment, we estimated the phase of gamma at which every spike occurred for each neuron, obtaining an average vector for which the mean resultant length (MRL) was calculated (see methods). This MRL is an estimative of the phase-locking strength for every neuron. The mean MRL to gamma was higher during searching compared to exploration in spatial strategies, with no differences between stages in non-spatial strategies (spatial: P < 0.0001; non-spatial: P = 0.306; Sidak’s multiple comparisons test; **Fig. 5b**). Then, we asked if hippocampal and parietal 4-Hz and theta oscillations synchronized neuronal spiking in the mPFC. As shown in examples of recordings in **Figs. 5c, e, g**, and **i**, prefrontal neurons fired aligned to the phase of hippocampal and parietal oscillations. We found that entrainment of prefrontal neurons by hippocampal 4-Hz was higher during searching compared to exploration stages during spatial, but not during non-spatial strategy (spatial: P < 0.0001; non-spatial: P = 0.824; Sidak’s multiple comparisons test; **Fig. 5d**). Similar results were found for parietal-4-Hz modulation of prefrontal firing (spatial: P < 0.0001; non-spatial: P = 0.995; Sidak’s multiple comparisons test; **Fig. 5h**). On the other hand, the modulation of prefrontal firing by hippocampal theta was higher during searching, irrespective of the strategy implemented (spatial: P < 0.0001; non-spatial: P < 0.0001; Sidak’s multiple comparisons test; **Fig. 5f**). Similar results were found for entrainment by parietal theta (spatial: P < 0.0001; non-spatial: P < 0.0001; Sidak’s multiple comparisons test; **Fig. 5j**).

**Figure 5.**
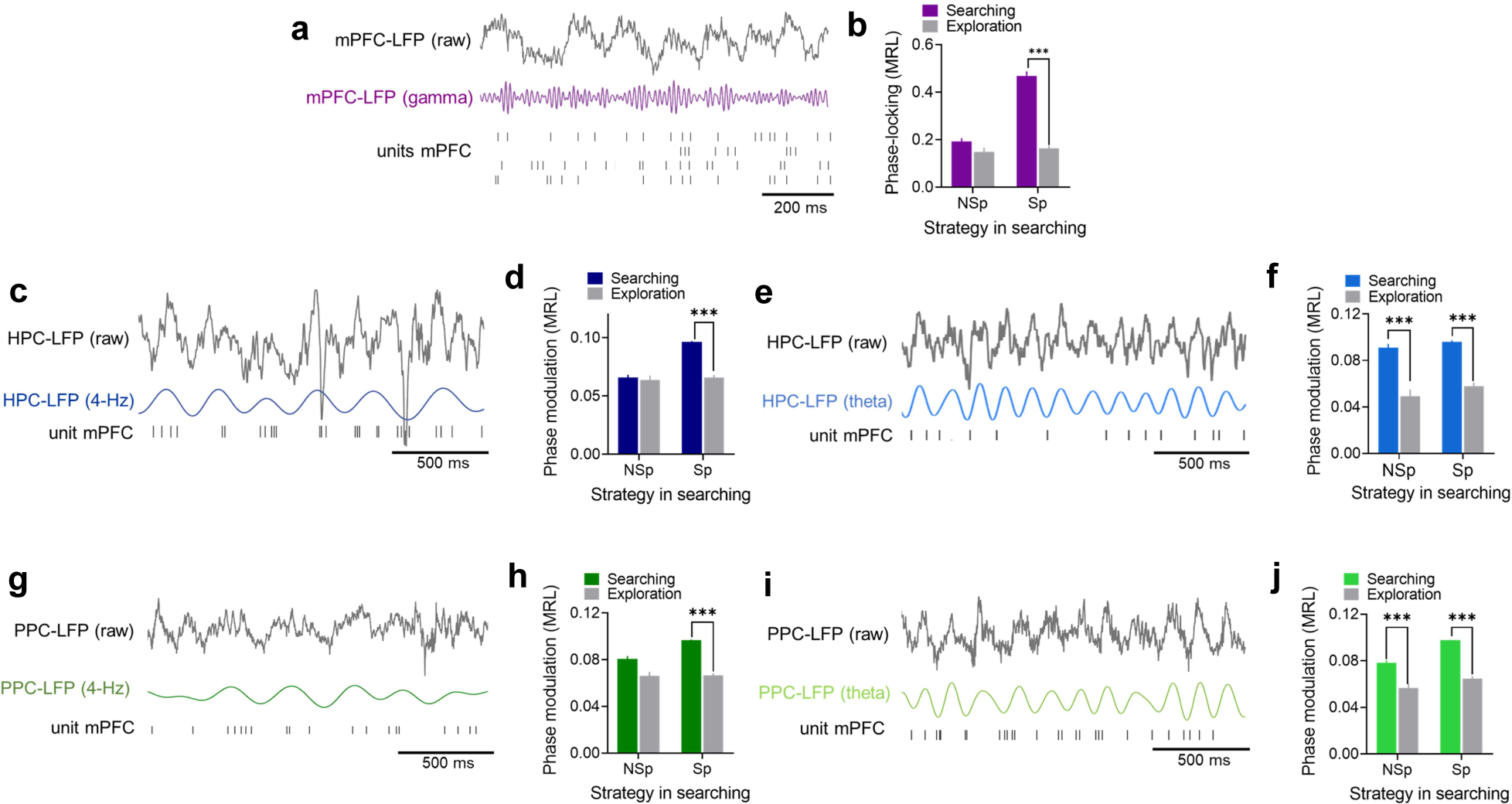
The firing of prefrontal neurons is modulated by local gamma and long-range hippocampal and parietal 4-Hz and theta oscillations. **a)** Example of prefrontal raw (black, upper) and gamma-filtered (purple, middle) LFP recording, and raster plot of spikes of four simultaneously recorded neurons from mPFC (lower). **b, d, f, h, j)** Bar chart of average phase-locking strength (measured as MRL) as function of navigation strategy and task stage for all neurons recorded from mPFC respect to local gamma oscillations **(b)**, hippocampal 4-Hz and theta (**d** and **f**, respectively), and parietal 4-Hz and theta oscillations (**h** and **j**, respectively). Data are presented as mean ± SEM. ***: P < 0.001; Sidak’s multiple comparisons test after two-way ANOVA. **c, g)** Raw (upper) and 4-Hz filtered (middle) LFP recording from the HPC **(c)** and PPC **(g)**, and spikes of a prefrontal neuron modulated by hippocampal **(c)** and parietal **(g)** 4-Hz oscillations. **e, i)** The same as **c** and **g**, but for prefrontal neurons modulated by hippocampal and parietal theta oscillations.

### 3.5. Reorganization of firing patterns in the mPFC during spatial learning

We finally evaluated if firing patterns in the mPFC were modulated according to task stages and navigation strategies. Given that the variability of the duration of task stages does not allow direct comparison of firing patterns between different training sessions, each stage was subdivided into an equal number of bins, and the normalized (z-scored) firing rate was calculated for each bin. Thus, it is possible to compare the dynamics of spiking relative to relevant behavioral events independently of the duration of each task stage. These data are shown in the color-coded raster plot in **Fig. 6a**, in which normalized firing was segregated according to firing in turn to stage transition (i.e., the first nose poke in the escape hole). From this figure, it can be seen that, when mice used spatial strategies in the searching stage, several neurons changed their firing pattern between stages. Indeed, prefrontal units showed more complex firing during searching when they used spatial strategies. Contrarily, when mice used non-spatial strategies during searching, firing remained relatively constant during the entire navigation session (**Fig. 6a**). This is most evident in the normalized peri-event time histogram of the lower panel in **Fig. 6a**. To quantify the differences in firing patterns between stages, we calculated the stage-associated firing selectivity index (FSI) for each neuron (see methods). Thus, the firing of each neuron was expressed as a value between -1 and 1 that estimates its dynamics between stages; positive values indicate that firing was higher in searching compared to exploration; negative values indicate the opposite; values near 0 indicate no changes between stages; and the absolute value indicates the magnitude of the variation. To evaluate if firing dynamics changed with strategy progression, we compared the distribution of FSI values between navigation strategies. As shown in **Fig. 6b**, the values of FSI showed a distribution toward more extreme values during spatial strategies compared with non-spatial strategies. Statistical analysis revealed significant differences in the distribution of FSI values between non-spatial and spatial strategies (P = 9,7 e-21; Kolmogorov-Smirnov test; **Fig. 6b**). To quantify the strength of these dynamics, we compared the magnitude of change in firing patterns (absolute FSI value) between strategies (**Fig. 6c**). We found that absolute FSI values were higher during spatial strategies compared to non-spatial (P < 0.0001; Mann-Whitney test; **Fig. 6c**), indicating that the prefrontal neuronal population increases their selectivity of firing in a given task stage during spatial strategies compared to non-spatial strategies.

**Figure 6.**
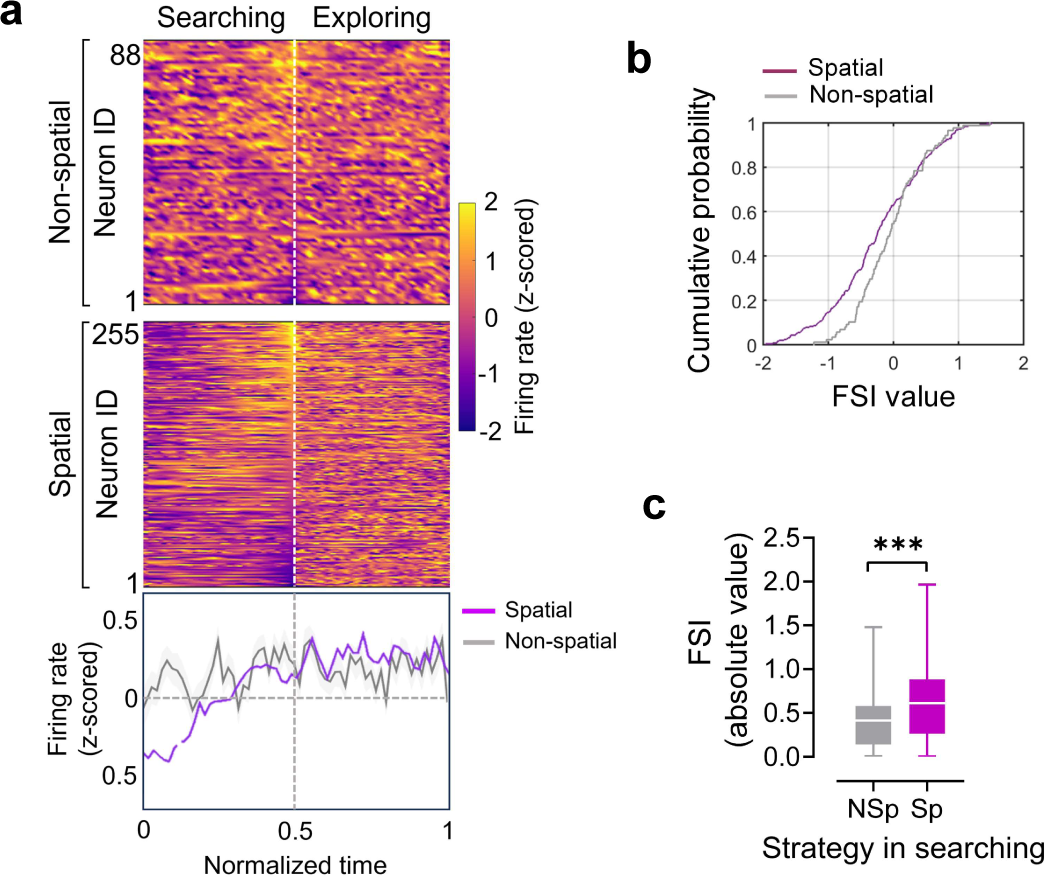
Dynamics of firing patterns of prefrontal neurons during spatial memory acquisition. **a)** Upper panel: color-coded normalized time and z-scored firing rate for all neurons recorded in the mPFC aligned to behaviorally relevant events during non-spatial (upper) and spatial (lower) strategies. Lower panel: average z-scored peri-event time histogram during non-spatial (grey) and spatial strategies (purple). **b)** Cumulative distribution of FSI for all neurons recorded in the mPFC during non-spatial and spatial strategies. **c)** Box plot comparing the absolute value of the FSI between non-spatial and spatial strategies for all neurons recorded in the mPFC. The middle, bottom, and top lines correspond to the median, lower, and upper quartiles, and the edges of the lower and upper whiskers correspond to the 5th and 95th percentiles. ***: P < 0.001; Mann-Whitney test.

## 4. DISCUSSION

In this study, we found that mice sequentially executed a searching and an explorative behavior along navigation trials, in which strategy progression was observed only in the searching stage. Interestingly, only when animals searched for the escape hole using spatial navigation strategies, 4-Hz and theta oscillations from the HPC and PPC synchronized gamma activity and firing patterns in the mPFC, which coincided with the highest incidence of gamma oscillations, the entrainment of firing patterns by these local gamma oscillations, and the task-related accommodation of firing patterns in the mPFC. These findings suggest that the prefrontal cortex is transiently recruited and large-scale coupled in a task-specific, learning-dependent manner.

Our behavioral analysis suggests that the searching stage was directed toward goal achievement (**Fig. 1e, g**). At the cognitive level, this requires the control and organization of behavior, a function supported by the mPFC (Eichenbaum, 2017; Fuster, 2008; Negrón-Oyarzo et al., 2024). Therefore, the increased prefrontal activity and synchronization observed during the searching stage may promote neural operations implicated in behavioral control for goal achievement. On the other hand, we found that, in agreement with current literature (Modirshanechi et al., 2023), exploratory behavior was not goal-directed (**Fig. 1e, g**). In a consistent manner, the reduced prefrontal synchronization observed during exploration may imply a reduced recruitment of the mPFC during this stage. Therefore, this evidence suggests that prefrontal activity synchronization is transiently implemented when goal-directed guide of behavior is required. Notably, prefrontal activity synchronization during searching was evidenced during the implementation of spatial navigation strategies, that were based on the modification of the navigation path (**Fig. 1f; supplementary Fig. 1l, m**) and strongly impacted behavioral performance (**supplementary Fig. 1n, o**). Spatial strategies occurred preferentially at the end of the training process (**Fig. 1g**), suggesting that the goal-directed accommodation of behavior, and the consequent recruitment of prefrontal activity, may require accumulated experience acquired throughout previous navigation trials (Negrón-Oyarzo et al., 2018; Ruediger et al., 2012). Present learning theories propose that a generalized internal map-like representation is gradually constructed during experience accumulation through the systematic organization, generalization, and extraction of commonalities from past similar episodes (Eichenbaum, 2017; Preston & Eichenbaum, 2013; Schlichting & Preston, 2015). Thus, this generalized map can be subsequently utilized to flexibly guide goal-directed behavior according to current conditions (Tang & Jadhav, 2022). Both the construction of the internal representation (Eichenbaum, 2017; Preston & Eichenbaum, 2013; Schlichting & Preston, 2015), as well as the implementation of efficient strategies during spatial learning are supported by the mPFC (de Bruin et al., 1994; Kesner et al., 1989; Kolb et al., 1994; Patai & Spiers, 2021). Thus, our data suggest that the prefrontal neural processing required for goal-directed accommodation of behavior may emerge once sufficient experience has been accumulated.

For the execution of efficient strategies, the mPFC may integrate several sets of information into the current neural operations. Space-based information and action-based self-motion information are required for spatial learning (Robinson et al., 2020; Shamash et al., 2023). These items of information are represented by neural spiking in the HPC and PPC, respectively (Alexander et al., 2022; Buzsáki & Moser, 2013; Orban et al., 2021; Whitlock, 2017), structures that are anatomically connected with the mPFC (Jay & Witter, 1991; Kolb & Walkey, 1987; Swanson, 1981; Zingg et al., 2014). Spatial information from the HPC is transmitted to the mPFC (Nishimura et al., 2021), which synchronizes with the firing of prefrontal neurons (Bota et al., 2021; Nardin et al., 2021; Tang et al., 2017). Interestingly, whereas hippocampal representation of space remains specific, prefrontal spatial maps generalize across environments (Tang et al., 2023). Considering that hippocampal spatial maps are formed in the first exposition to the environment (Frank et al., 2004; Hill, 1978), it can be suggested that spatial information requires some “executive” processing before it can be used to guide behavior. Similarly, whereas neurons in the PPC represent accumulated evidence, prefrontal neurons encode categorical features in memory-guided tasks (Hanks et al., 2015; Scott et al., 2017). Therefore, the transference of information from the HPC and PPC into the mPFC may contribute to the formation of generalized categorical maps in the mPFC as required to guide behavior during learning.

Large-scale transference and integration of information across brain regions can be mechanistically promoted by the entrainment of local high-frequency oscillations by low-frequency oscillations (Canolty & Knight, 2010; Hyafil et al., 2015; Lisman & Jensen, 2013). Interestingly, we found that hippocampal and parietal 4-Hz and theta oscillations entrained gamma oscillations in the mPFC (**Fig. 3**). This entrainment was highest when animals executed spatial strategy in the searching stage (**Fig. 3b, c**). The CFC analysis also revealed that both 4-Hz and theta oscillations entrained gamma oscillations at a similar central frequency (near 80 Hz; **Fig. 4b, c**). Also, gamma oscillations increased their incidence at an analogous central frequency, specifically in the mPFC when animals executed spatial strategies during the searching stage (**Fig. 4b, c**). This suggests that 4-Hz and theta oscillations modulated gamma oscillations as a common neurophysiological process in the mPFC. Furthermore, spike timing of prefrontal neurons was modulated by local gamma oscillations, as well as by hippocampal and parietal 4-Hz and theta oscillation modulation, which also increased when animals executed spatial strategy during the searching stage (**Fig. 5**). Gamma oscillations emerge as a result of the excitatory and inhibitory interaction between local neural populations (Buzsáki & Wang, 2012). Entrainment of prefrontal neurons by long-range oscillations modulates the instantaneous spiking probability (Fell & Axmacher, 2011), facilitating the interaction between local neurons and thus promoting the emergence of gamma oscillations. Hence, via the entrainment of neuronal spiking, large-scale CFC may promote the interaction between prefrontal neurons, increasing CFC and gamma incidence in the mPFC. Considering that gamma oscillations promote neural operations for cortical computations, as associative binding of distributed representations (Fernandez-Ruiz et al., 2023; Fries, 2009; Griffiths & Jensen, 2023), gamma oscillations may promote the integration of information for cognitive control during goal-directed spatial learning.

Our analysis of the CFC revealed that 4-Hz and theta oscillations are two distinct potential approaches to long-range entrain prefrontal gamma in the mPFC. These oscillations do not coexist during spatial learning (**Fig. 2c** and **supplementary Fig. 2a**), and they differ in their circuital origin in the mPFC, HPC and PPC (F. Jung et al., 2022; Karalis & Sirota, 2022; Yanovsky et al., 2014). In awake rodents, theta emerges mostly during locomotion (**Fig. 2d**); therefore, this rhythm is the main global integrator when animals move in the environment (Colgin, 2013). However, cognitive processing is also implemented in the absence of locomotion and, consequently, in the absence of theta oscillations. This occurs, for example, during deliberative processes (Redish, 2016). Therefore, the exclusive role of theta as a long-range integrator seems insufficient. Considering that 4-Hz oscillation emerges widely in the brain in the absence of locomotion (**Fig. 2d**; (Biskamp et al., 2017)), 4-Hz oscillation may work as a global integrator of neural operations when locomotion is absent (Folschweiller & Sauer, 2021; Heck et al., 2019). Interestingly, mPFC, HPC and PPC are apparently simultaneously recruited by either 4-Hz or theta oscillations (**Fig. 2b**). Evidence supporting widespread brain synchronization by theta or 4-Hz oscillations has been previously reported (Karalis & Sirota, 2022; Tort et al., 2018; Zhong et al., 2017). Thus, the mPFC, HPC and PPC may be coupled in two main large-scale configurations depending on locomotion: one dominated by the 4-Hz during moments of immobility, and the other dominated by theta oscillations during locomotion.

Finally, we found that prefrontal neurons modified their firing between stages according to navigation strategy. Stage-associated FSI values were more extreme and had higher absolute value during spatial strategies **(Fig. 6b, c**), indicating that neurons in the mPFC increased the specificity of firing according to navigation efficiency. Indeed, the firing pattern of a large proportion of prefrontal neurons seems less dispersed during searching compared with the exploration stage in spatial strategies (**Fig. 6a**). This implies that firing patterns were adjusted during the searching stage depending on the strategy used. Indeed, it has been previously found that prefrontal firing patterns signal the approach to the goal when mice use spatial strategy (Negrón-Oyarzo et al., 2018). Therefore, firing patterns may be more specific to particular behavioral events during searching than during exploration. This issue will be approached in a future and deeper analysis.

Overall, our results provide evidence for the large-scale coordination of the mPFC with the HPC and PPC as a putative neural mechanism supporting spatial learning. This mechanism was primarily implemented when mice organized their trajectory in an efficient goal-directed manner and was executed after sufficient experience had been acquired. It involved theta and 4-Hz oscillations that emerged differentially according to locomotor activity. These oscillations coordinated gamma oscillations and neuronal firing in the mPFC, which coincided with the increase of gamma oscillations and the task-related arrangement of firing patterns in the mPFC.

## 5. MATERIALS AND METHODS

### Animals

Adult male C57BL/6j mice (n = 12; age: 60–90 days) were used in this study. Mice were housed in a 12-hour light/dark cycle in a temperature- and humidity-controlled room (22 ± 2 °C) with ad libitum access to food and water. All experimental procedures related to animal experimentation were approved by the Institutional Animal Ethics Committee of the Universidad de Valparaíso (protocol code: BEA178-22). Efforts were made to minimize the number of animals used and their suffering.

### Fabrication of microelectrode arrays (MEA)

Custom-made MEA carrying tungsten wire microelectrodes (SML coated; 50 µm diameter, California Fine Wire) were assembled for simultaneous LFP and single units recording from the mPFC, CA1 area of the HPC, and the PPC. Microelectrode wires were cut off with iris scissors (World Precision Instruments, Sarasota Instruments). The final impedance of each wire (200–500 kΩ) was measured in a saline solution at 1 kHz. Microelectrode arrays were composed of two bundles of seven stainless-steel cannula each (30 G; Components Supply Co., FL, USA), one directed to mPFC and the other to HPC/PPC. Six single wires were inserted in each bundle, and the length was adjusted to target the mPFC (length: 1.5 mm), the HPC (length: 1.5 mm), and the PPC (length: 0.5 mm). Each microelectrode wire was fixed to the bundle with cyanoacrylate and connected to a 16-channel interface board assembled with an Omnetics connector (Neurotek, Toronto, ON, Canada).

### MEA implantation surgery

Animals were anesthetized with isoflurane (3% isoflurane with 0.8% O2) before being placed in a stereotaxic frame. Anesthesia was maintained until the surgery was finished (1–2% isoflurane with 0.8% O2). After incision in the scalp, two craniotomies were drilled in the right hemisphere at stereotaxic coordinates targeting the mPFC (1.94 mm AP, -0.25 mm ML, from Bregma; (George Paxinos, 2001)) and region CA1 of the HPC (-1.94 mm AP, - 1.5 mm ML, from Bregma). Stereotaxic coordinates targeting PPC were the same as for HPC. The dura was removed, and the electrodes were slowly lowered and inserted into the cortical surface. Two ground wires were attached to skull screws. Once in position, the MEA and the ground screws were affixed to the skull with dental acrylic. After surgery, animals were maintained in individual cages in a temperature- and humidity-controlled room (22 ± 2 °C) with food and water ad libitum and were supplied with a subcutaneous dose of analgesics (Ketoprofen, 5 mg/kg/day) and antibiotics (Enrofloxacine, 5 mg/kg/day) during the 5 days after surgery. Mice were allowed to recover for at least one week after surgery before beginning behavioral and recording experiments. During recovery, weight and general health were monitored daily.

### Behavioral and electrophysiological recording procedures

After one week of recovery from surgery, mice were habituated and placed in the behavioral room for 15 minutes, and then they were trained for the the the the spatial reference memory task in the Barnes maze (Barnes, 1979). The maze consisted of a white circular platform of 70 cm in diameter elevated at 70 cm from the floor with 16 equally spaced holes (9 cm diameter) along the perimeter and located at 2 cm from the edge of the platform. Visual cues were located on the walls of the room. Under one of the holes was located a black plexiglass escape box (17 x 13 x 7 cm) that allowed the mice to enter with the implanted MEA. The location of the escape- box was consistent for a given mouse but randomized across the mice group. The spatial location of the target was unchanged in respect to the distal visual room cues. The maze was illuminated with two incandescent lights to yield a light level of approximately 400 lux impinging on the circular platform. To evaluate the acquisition of spatial memory, four navigation trials per day with an inter-trial interval of 15 minutes during 5 consecutive days were realized. In each acquisition trial, the interface board of the MEA was connected to the amplifier board (RHD2000 evaluation system; Intan Tech, CA, USA) via a 16-channel head-stage (model RHD2132 Intan Tech, CA, USA). Then the recording system was turned on, and the mouse was placed in the start box in the center of the maze for 1 min with the room lights turned off (start stage). After time had elapsed, the start-box was removed, the room lights turned on, and the mouse was free to explore the maze (navigation stage). The session ended when the mouse entered the escape box or after 3 minutes elapsed. Once the mouse entered the escape, the lights were turned off, and the mouse was left to remain in the escape box for 1 min (goal stage). If the mouse did not enter the escape box within 3 minutes, the experimenter guided the mouse to the escape. When the trial was completed, the electrophysiological recording system was turned off, the mouse was disconnected from the electrophysiological recording system and returned to the home cage, and the maze was cleaned with ethanol 70% to prevent a bias based on olfactory or proximal cues within the maze. During the complete trial the animal behavior was tracked and recorded with a webcam (acquisition at 30 fps and 640 × 480 pixels; camera model C920; Logitech Co.) located 1 mt above the maze and controlled with VirtualDub software.

### Histology

After the complete experiment was finished, mice were anesthetized with 3% isoflurane, and the location of each microelectrode wire was marked by passing a small amount of current through each electrode (50 µA by 10s). Two days after this protocol, mice were anesthetized with isoflurane (3%) and then transcardially perfused with ∼40ml of phosphate-buffered saline (PBS, pH = 7.4), followed by ∼50ml of paraformaldehyde (PFA) in PBS. The brain was removed and then stored in PFA with PBS overnight. PFA was washed with a solution of PBS containing 30% saccharose, and brains were stored in PBS with 30% saccharose until the cut session. Coronal brain slices (50 µm) from HPC/PPC and mPFC were obtained from a cryostat-microtome (Zhejiang Jinhua Kedi Instrumental Equipment Co., LTD) and stored in PBS containing 0.01% sodium azide. For visualization of the electrolytic lesions, slices were stained with Nissl-staining, and pictures were acquired with a microscope (Nikon).

### Behavioral analysis

#### Behavioral performance

Behavioral performance was analyzed by measuring the escape latency (the time to enter the escape hole), nose-poke latency (time elapsed for the first contact of the mouse nose with the escape hole), number of errors (nose-pokes in non-escapes holes), distance covered and path efficiency to the first nosepoke in the escape hole. To assess locomotor activity, the mean and maximum speed in all trials were measured. All these behavioral parameters were analyzed and measured with ANY-maze software (Stoelting Co.). Also, ANY-maze software provided for every trial a file containing instantaneous spatial coordinates of the animal’s trajectory and instantaneous speed. These were used to construct occupancy plots and for the detection of episodes of mobility (running speed > 4 cm/sec for at least 2 sec) and immobility (running speed < 4 cm/sec for at least 2 sec) during navigation. Both analyses were performed in MATLAB software with custom-made scripts.

#### Identification of task stages

For the identification of relevant task moments and comparison of electrophysiological parameters among them, each navigation trial was separated into two stages: searching, from the start of navigation to the first nose poke in the escape hole; and exploration, from the first nose poke in the escape hole to entering the escape hole. *Identification of navigation strategies:* navigation strategies were defined and analyzed with the Barnes Maze Unbiased Strategy Classification Tool (BUNS)(Illouz et al., 2016) implemented in MATLAB software. Coordinates of animal trajectory obtained from ANY-maze software were entered in MATLAB by using the support vectorial machine (SVM) and converted into cartesian coordinates to characterize animal trajectory features until the first contact with the escape hole. Data were adjusted at BUNS detector requirements and then the strategies were classified according to the following definitions: random, when the animal navigates randomly in the maze until that it finds the escape-hole; serial, when the animal navigates of hole to hole (adjacent holes) until it finds the escape-hole; focused search, when the animal realizes a located scan of the escape-hole quadrant; long correction, when the animal navigates to erroneous hole and, then it adjusts the navigation trajectory to directly find the hole escape; corrected, when the animal slightly modifies its trajectory (go to the next hole, before the escape-hole); direct, when the animal uses the shortest path to find the escape-hole. All the results obtained from the strategy detector were compared with a visual analysis recording to determine possible errors. BUNS also provides a cognitive score according to the strategy (direct = 1 point; corrected = 0.75 points; long correction = 0.5 points; focused search = 0.5 points; serial = 0.25 points; and random = 0 points).

#### Path sparsity

Path sparsity, which represents the proportion of arena the animal occupied to reach the goal, was based on the method described by Jung et al. (M. W. Jung et al., 1994). Briefly, to parametrize the sparsity for firing rate, trajectory sparsity was calculated as follow:

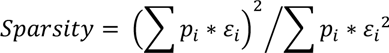

where *p_i_* is the probability of the animal being in the *i* th bin (occupancy time in the ith bin/total recording time) and *ε_i_* is the total time, the animal visits the *i* th bin.

### Electrophysiological analysis

#### LFP spectral power and instant energy

For analysis of the spectral power of oscillatory activity, electrophysiological recordings were down-sampled to 1000 Hz, and bandpass-filtered at 0.1–120 Hz. Power spectral density (PSD) was computed using multi-taper Hamming analysis using the Chronux toolbox (http://www.chronux.org; (Mitra & Pesaran, 1999)). For PSD analysis, field potentials were divided into 4000 ms segments with 400 ms overlap and a time-bandwidth product (TW) of 5-9 tapers. For the estimation of spectral power during mobility and immobility episodes, the down-sampled LFP recordings were filtered at 4-Hz or theta frequency bands using the Hilbert transform, and the resultant envelope (i.e., instant energy) was rectified, smoothed, and z-scored (Belitski et al., 2010).

#### Gamma-event detection

For gamma event detection we used the method described by Logothetis et al (Logothetis et al., 2012) with some variations from Csicsvari et al. (Csicsvari et al., 2003). Briefly, LFP was band-pass filtered (60–100 Hz) using a zero-phase shift noncausal finite impulse filter with a 0.5 Hz roll-off. Next, the signal was rectified and low-pass filtered at 20 Hz with a 4th-order Butterworth filter. This procedure yields a smooth envelope of the filtered signal, which is then z-score normalized using the mean and standard deviation (SD) of the whole signal. Epochs during which the normalized signal exceeded a 2.0 SD threshold were considered potential gamma events. The first point before threshold that reached 1 SD was considered the onset, and the first one after threshold to reach 1 SD as the end of events. The difference between the onset and end of events was used to estimate the gamma event duration. We introduced a 50-ms-refractory window to prevent double detections. In order to precisely determine the mean frequency, amplitude, and duration of each event, we performed a spectral analysis using Morlet complex wavelets of 7 cycles.

#### Cross-frequency coupling analysis

For CFC analyses was used the ModIndex toolbox (Tort et al., 2010). CFC was computed using the Modulation Index (MI). The MI is able to detect the strength of the phase-amplitude coupling between two frequency ranges of interest: the “phase-modulating” (theta and 4-Hz) and “amplitude-modulated” (gamma) frequency bands. First, LFP was filtered at the two frequency ranges under analysis. Next, the phase and the amplitude time series were calculated from the filtered signals by using the Hilbert transform. Specifically, the time series of the phases were obtained from the standard Hilbert transform, and to extract the amplitude envelope, we also applied the Hilbert transform. Then, the composite time series was constructed, which gave us the amplitude of the “amplitude-modulated” oscillation at each phase of the “phase-modulating” oscillation. The phases were binned, and the mean amplitude over each phase bin was calculated. The existence of phase-amplitude coupling is characterized by a deviation of the amplitude distribution P from the uniform distribution U in a phase-amplitude plot (if there is no phase-amplitude coupling, the amplitude distribution P over the phase bins is uniform). A measure that quantifies the deviation of P from the uniform distribution was determined by an adaptation of the Kullback-Leibler (KL) distance (DKL), which is widely used in statistics to infer the amount of difference between two distributions. The adaptation was to make the distribution distance measure assume values between 0 and 1. The MI is therefore a constant time, the KL distance of P from the uniform distribution (U). Thus, the higher the MI, the greater the phase-amplitude coupling between the frequencies and areas of interest. It is represented in the following equation:

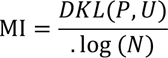

#### Spike sorting

Spike sorting was performed offline using Wave-clus, a MATLAB based super paramagnetic clustering method for spike sorting (Quian Quiroga & Nadasdy, 2004). To detect spikes, broadband recordings sampled at 20 kHz were filtered at 500–5000 Hz, and events with amplitudes reaching 4–50 standard deviations over the mean were considered candidate spikes. For each recording file, single channels from each single electrode were identified, and a single file was generated per channel. The spike features considered for clustering included peak amplitude (the maximum height of the waveform of each channel of each spike), timestamp (the time of occurrence of each spike), spike width (the duration in each channel of the spike), energy (the energy contained within the waveform of each channel of the spike), valley (the maximum depth of the waveform of each channel of the spike), and wave-PC (the contribution to the waveform due to the principal component). A file for every single unit, including timestamps of spikes, was generated and used in the subsequent analyses on MATLAB. To confirm the correct sorting of single units, the time stamps of every single unit were aligned with their 500–5000 Hz filtered LFP recording, and the concordance of timestamps with spikes was visually established.

#### Phase-locking analysis

Phase-locking analysis was computed using the Matlab toolbox CircStats (http://philippberens.wordpress.com/code/circstats/; (Berens, 2009)). Briefly, LFP traces were bandpass filtered at 4-Hz, theta or gamma range (2–5 Hz, 6–12 Hz, and 60–100 Hz, respectively; zero phase shift non-causal finite impulse filter with 0.5 Hz roll-off). Phase locking was quantified as the circular concentration of the resulting phase distribution, which was defined as mean resultant length (MRL = (n/Z)0.5)(Adhikari et al., 2011). The statistical significance of phase locking was assessed using the Rayleigh test for circular uniformity. To avoid bias, we only considered neurons with >50 recorded spikes.

#### Firing selectivity index analysis

For the calculation of the stage-associated firing selectivity index, we used the following expression:

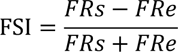

in which, for any given single unit, FRs is the mean firing rate during searching, and FRe is the mean firing rate during exploration. Values may range from -1 to 1. Positive values indicate that firing was higher in searching compared to exploration; negative values indicate the opposite; values near 0 indicate no changes between stages; and the absolute (abs(FSI)) value indicates the strength of the variation.

### Statistical analysis

Comparisons between the means of quantitatively normally distributed variables in two groups were analyzed using the t-student test. Comparisons between the means of quantitatively non-normally distributed variables in two groups were analyzed using the Mann-Whitney test. Comparisons between the means of quantitatively normally distributed variables of more than two groups (as most behavioral variables) were analyzed using one-way ANOVA followed by Tukey’s multiple comparisons. Comparisons between the means of quantitative variables across two categorical independent variables (for example, MI or MRL between stages and navigation strategies) were performed using a two-way ANOVA followed by Sidak’s multiple comparisons test. Linear correlations between parameters were analyzed by the Pearson correlation test. Distributions (SI values) were compared using the Kolmogorov-Smirnov test. Statistical analysis was performed with GraphPad Prism or MATLAB (The Mathworks Inc.) software. Significance differences were accepted at P < 0.05.

## 6. ACKNOLEDMENTS

This work was supported by ANID Chile, projects ANILLO ACT210053 to I.N-O., P.F. and W.E-D.; FONDECYT Iniciación N°11160251 to I.N-O and Postdoctorado N°3230400 to FG; and Beca de Doctorado Nacional to L.C-V. I.N-O acknowledges the support of the Universidad de Valparaíso: INICI-UV (UVA20993), and Centro de Neurobiología y Fisiopatología Integrativa (DIUV-CI 01/2006). We thank Dr. Pablo Moya, Dra. Ana María Cárdenas, Dr. Alexies Dagnino, and Dr. Ramón Sotomayor for sharing physical space and equipment for histological experiments.

## 7. AUTHOR CONTRIBUTION

I.N-O. designed the experiments. F.G., L.C-V, and I.N-O. carried out the experiments. F.G., M-J.T. and I.N-O. analyzed the data. I.N-O. and N.E. developed analytical scripts. I.N-O. W.E-D and P.F. interpreted the data. I.N-O. wrote the manuscript. All authors discussed and commented on the manuscript.

## Competing financial interests

The authors declare no competing financial interests.

## SUPPLEMENTARY FIGURES

**Supplementary figure 1.**
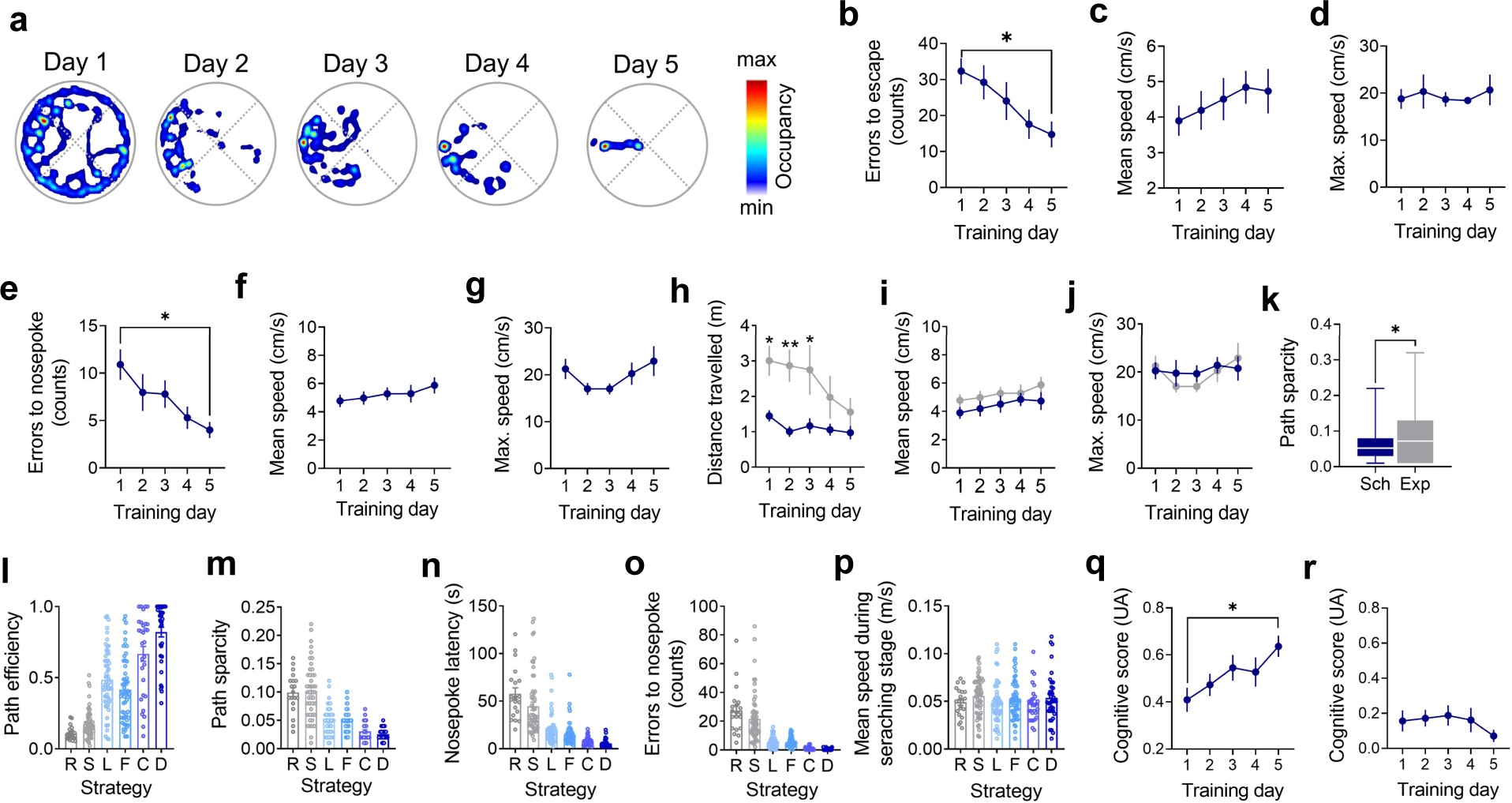
**a)** Example of color-coded occupancy plots in the Barnes maze across training days. **b-g)** Comparison of errors to escape **(b)**, mean speed **(c)**, maximum speed **(d)**, errors to nose poke **(e)**, mean speed **(f)**, and maximum speed **(g)** across training days. Data are presented as mean ± SEM. *: P < 0.05; Tukey’s multiple comparisons test after one-way ANOVA. **h-j)** Comparison of distance traveled **(h)**, mean speed **(i)**, maximum speed **(j)**, between searching and exploration stages. Data are presented as mean ± SEM. **: P < 0.01; *: P < 0.05; Sidak’s multiple comparisons test after two-way ANOVA. **k)** Box plot comparing path sparsity between searching and exploration stages. *: P < 0.05; t-student test. **l-p)** Bar chart comparing path efficiency **(l)**, path sparsity **(m)**, nose poke latency **(n)**, error to nosepoke **(o)**, and mean speed **(p)** across goal-searching strategies during the searching stage. Each point represents a single trial session of training in the Barnes maze. **q-r)** Comparison of cognitive score during the searching **(q)** and exploration stages **(r)** across training days. Data are presented as mean ± SEM. *: P < 0.05; Tukey’s multiple comparisons test after one-way ANOVA.

**Supplementary figure 2.**
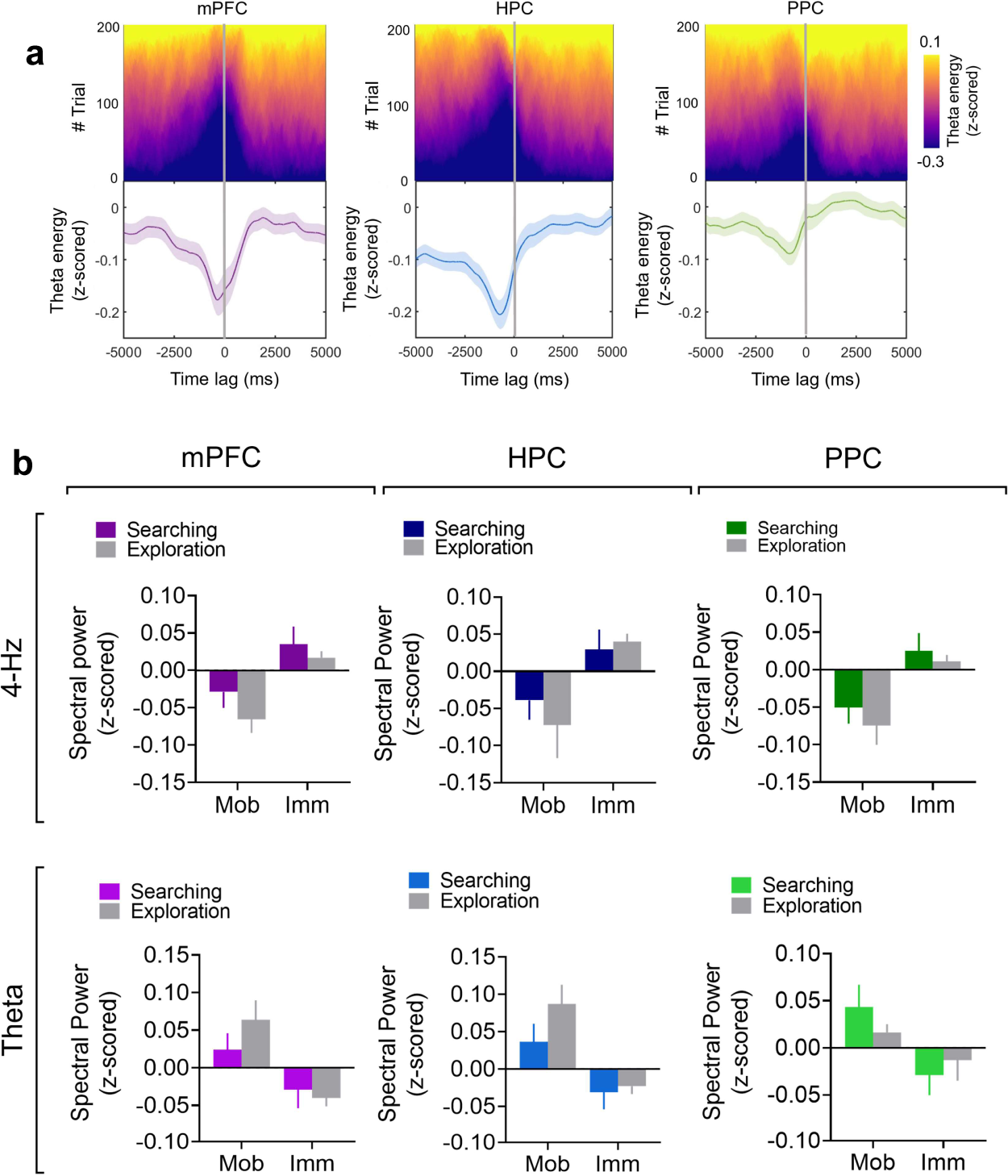
**a)** Upper panel: colorplot of z-scored energy of theta oscillation in turn to the of peaks 4-Hz oscillation in the mPFC (left), HPC (middle), and PPC (right) for all training sessions. Lower panel: average theta z-scored energy in turn to 4-Hz peaks. Data are presented as mean (solid line) ± SEM (shaded area). **b)** Upper panel: bar chart of mean z-scored spectral power of 4-Hz in the mPFC (left), HPC (middle), and PPC (right) with respect to mobility and immobility episodes in searching and exploration stages during training sessions. Lower panel: same as upper panel but for z-scored spectral power of theta oscillation. Data are presented as mean ± SEM.

**Supplementary figure 3.**
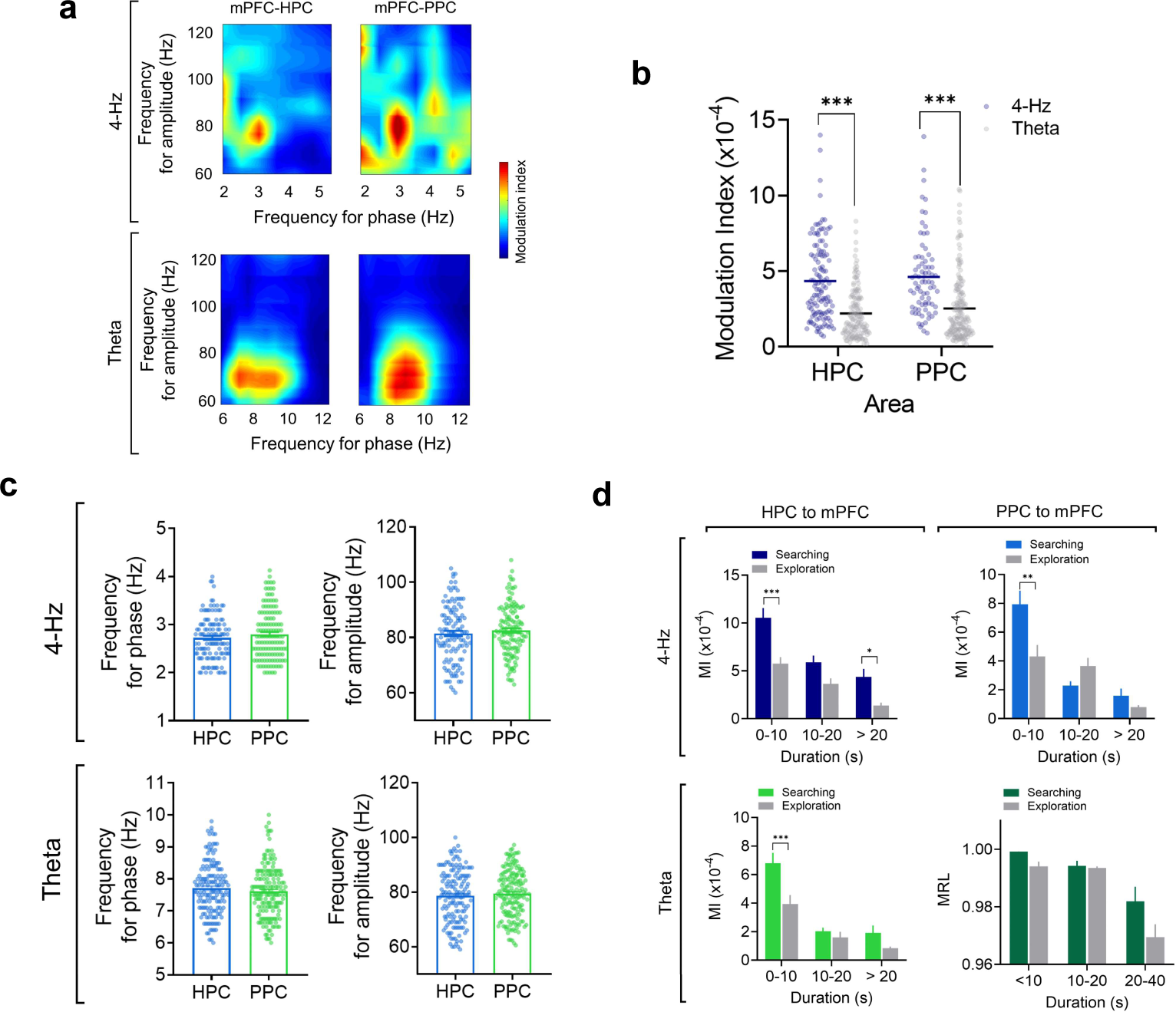
**a)** Examples of color coded comodulograms of CFC of prefrontal gamma respect to 4-Hz (upper) and theta (lower) oscillations from HPC and PPC. **b)** Raster plot comparing the Modulation Index of CFC of prefrontal gamma respect to 4-Hz and theta oscillations from HPC and PPC. Each point represents a single trial session of training in the Barnes maze. Solid line represents the mean. ***: P < 0.001; Sidak’s multiple comparisons test after two-way ANOVA. **c)** Bar chart and raster plot comparing the frequency for low-frequency phase (left) and the frequency for gamma amplitude of CFC of prefrontal gamma respect to 4-Hz (upper) and theta oscillations (lower) from HPC and PPC. Each point represents a single trial session of training in the Barnes maze. **b)** Bar chart comparing the modulation index of CFC between searching and exploration stages across sessions of similar duration for HPC to PFC (left) and PPC to PFC (right) and for 4-Hz (upper) and theta oscillations (lower). ***: P<0.001; Tukey’s multiple comparisons test after one-way ANOVA.

## Notes

### Competing Interest Statement

The authors have declared no competing interest.

